# Subtle, Seasonal and Potentially Important: Are Phytotoxic Foliage Leachates Overlooked in Plant Invasion Studies?

**DOI:** 10.1101/566976

**Authors:** Gruffydd Jones, Max Tomlinson, Rhys Owen, John Scullion, Ana Winters, Tom Jenkins, John Ratcliffe, Dylan Gwynn-Jones

**Affiliations:** IBERS, Aberystwyth University, Penglais Campus, Aberystwyth, Ceredigion. SY23 3DA, UK; Snowdonia National Park Authority, National Park Office, Penrhyndeudraeth, Gwynedd. LL48 6LF, UK; Forest Research, Thoday Building, Deiniol Road, Bangor, Gwynedd. LL57 2UW, UK; Natural Resources Wales, Maes y Ffynnon, Penrhosgarnedd, Bangor, Gwynedd. LL57 4DE, UK

**Keywords:** Phenolics, invasive, secondary metabolites, seasonal, inhibitory

## Abstract

Invasive species pose a threat to global biodiversity. Recently there has been a growth in research into the chemical aspect of invasions, with some plants reported to introduce compounds to the soil which impair the growth of competing native plants. One such species is *Rhododendron ponticum*, which has become a damaging invasive shrub in Britain. Here we investigate whether water-soluble compounds introduced from *R. ponticum* leaves in Ceredigion, Wales, suppressed seed germination, and whether this varied temporally. Leachates from leaf material collected monthly (March to July) were analysed for their chemical composition by liquid chromatography-mass spectrometry and *Lactuca sotiva* bioassays. In total, 19 compounds were identified, displaying both qualitative and quantitative seasonal variation. Total concentrations peaked in June, however *L sativa* germination inhibition was only found in leachates from April and May (15% and 17% respectively). The study provides evidence that phytotoxic chemical introduction from *R. ponticum* leaves is seasonal and most bioactive at a time of year when the seeds of other species germinate. We propose this is one of several mechanisms including shading, lowering of soil pH and the deposition of thick litter, that this species exploits when in competition with other species. We argue that for studies on the phytotoxicity of invasive plants, there is a need to consider temporal variation in the rate and quality of bioactive chemicals produced and released, local precipitation rates, persistence of chemicals in the soil plus synergistic interactions between individual chemicals.

## INTRODUCTION

Invasive species are increasingly damaging habitats worldwide, driven by the increased movement of goods and people due to globalisation (Perrings et al. 2005). The ecological factors behind plant invasions are well studied. Recent years however have seen a surge in interest in the chemistry of plant invasions, partly prompted by the “novel weapons hypothesis” proposed by Callaway and Aschehoug (2000). Many species are reported to possess such chemical traits, for example *Rhododendron ponticum* L., which has become a problematic invasive shrub in the UK following its introduction from Spain during the 18^th^ century (Cross 1975). In addition to its suitability to the British climate, some studies suggest *R. ponticum* also alters soil conditions by introducing bioactive secondary metabolites to the soil which impair the growth of competing native plants. Rotherham and Read (1988) found *Festuca ovina* L. (sheep’s fescue) root elongation was inhibited when growing in *R. ponticum* soil, an effect which was not alleviated by the addition of nutrients or water. This suggestion of phytotoxic compounds is supported by laboratory bioassays on litter leachates from another *Rhododendron* sp. (Nilsen et al. 1999).

Our investigation aimed to quantitatively and qualitatively study the seasonality of phenolic compounds in *R. ponticum* foliar leachates by liquid chromatography-mass spectrometry (LC-MS^n^), and the effect of these compounds on *Lactuca sativo* L. germination. It was hypothesised that these compounds are delivered via leachates from *R. ponticum* canopies and fallen leaf litter, with concentrations highest in spring (March, April and May) when the majority of seeds from other species germinate. Leading on from this, it was also hypothesised that leaf leachates would be most inhibitory to L *sativa* during these spring months, due to the reported phytotoxicity of some phenolic compounds.

## METHODS AND MATERIALS

### Site description

Sampling was conducted at a mixed woodland called Pwll Crwn (OSGB36 grid reference: SN623283), Ceredigion, Wales. This site was cleared of *R. ponticum* between May and July 2013 but plants had regenerated by March 2016. Samples were taken from 0.5 m stems regenerating from previously cut stumps.

### Leaf collection, phenolic compound analysis and bioactivity

Leaf leachates were collected monthly (March to July) from five independent randomly selected plants. Leaf extracts were used as opposed to soil extracts as it allowed the study to focus purely on the effect of the compounds derived from *R. ponticum;* soil extracts would contain compounds from other plants and microbes, as well as vary seasonally in terms of chemical properties. Separate plants were sampled each month to avoid any effects of leaf removal on subsequent secondary metabolite content. Extracts containing 3 g of sample tissue placed in 15 mL distilled water (1:5) were prepared using the method described in Nilsen et al. (1999). After storing at 4°C for 24 hours, 500 mg Sep-Pak C_18_ 3 cc Vac RC cartridges (Waters Ltd, Wexford, Ireland) were used to partially purify the extracts prior to LC-MS^n^ analysis. Following this, the extracts were evaporated to dryness with a vacuum centrifuge (Jouan RC1022, Nantes, France), before resuspending in 200 μL70% methanol.

The processed extracts were analysed by LC-MS^n^ on a Thermo-Finnigan LC-MS system (Thermo Electron Corp., Waltham, USA), consisting of a Finnigan Surveyor PDA Plus Detector, a Finnigan LTQ linear ion trap with ESI source and a 4 μm C_18_ Waters Nova-Pak column (3.9 mm x 100 mm). A constant temperature of 5°C was maintained for the autosampler tray, and 30°C for the column. The injection volume was 10 μL. The column was equilibrated with 95% of solvent A (water/formic acid; 100:0.1, v/v) and 5% solvent B (methanol/formic acid; 100:0.1, v/v). The percentage of solvent B increased linearly to 60% over 65 minutes, at a flow rate of 1 mL/min.

MS^2^ fragmentation data were used to identify compounds in negative ion mode. Compound concentrations were calculated using standard solutions of known volume for each of the chemical groups identified where available or the aglycone where compounds were not commercially available. Unknown compounds were quantified based on relative abundance.

Following on from these chemical analyses, *L sativa* (var. Buttercrunch) germination bioassays were conducted to investigate the impact these leachates have on plant growth, based on the method described by Nielsen et al. (1999). This species was selected as it is a standardised indicator species commonly used in studies on phytotoxic compounds. 20 seeds were evenly spaced on moistened (with 2 mL extract) filter paper in Petri dishes (55 mm); controls had the same volume of distilled water. Mean germination percentage (plus root and shoot length) per plate was recorded after incubation at 20 °C for seven days.

### Statistical analyses

Statistical analyses were conducted in SPSS (23.0, IBM, USA) using multiple regression and one-way ANOVA followed by the LSD post-hoc test (P < 0.05). Mean germination percentage was arcsine transformed prior to statistical analysis, whilst the concentration data were log transformed prior to multiple regression analysis.

## RESULTS

In total, 19 different phenolic compounds were detected, with all but two being identified (Table S2). Most compounds were present across all five months (March to July) with total concentration peaking in June (Fig. 1 and Table 2S) before decreasing significantly to July (P < 0.001). Whilst total phenolic concentration was highest in June, inhibition of *L sativa* germination was highest in April and May (P < 0.001) (Fig. 1). Of the 19 compounds detected in the leachates, only quercetin was found to be significantly more concentrated during a month where germination inhibition was observed; its concentration was higher in May compared to March and April (P < 0.05), whilst it was not present during June or July (Table 2S). A multiple regression analysis showed none of the 19 compounds had a significant relationship with germination (Table 1S), with only quercetin showing a near-significant negative relationship (P = 0.061).

**Figure 1:**
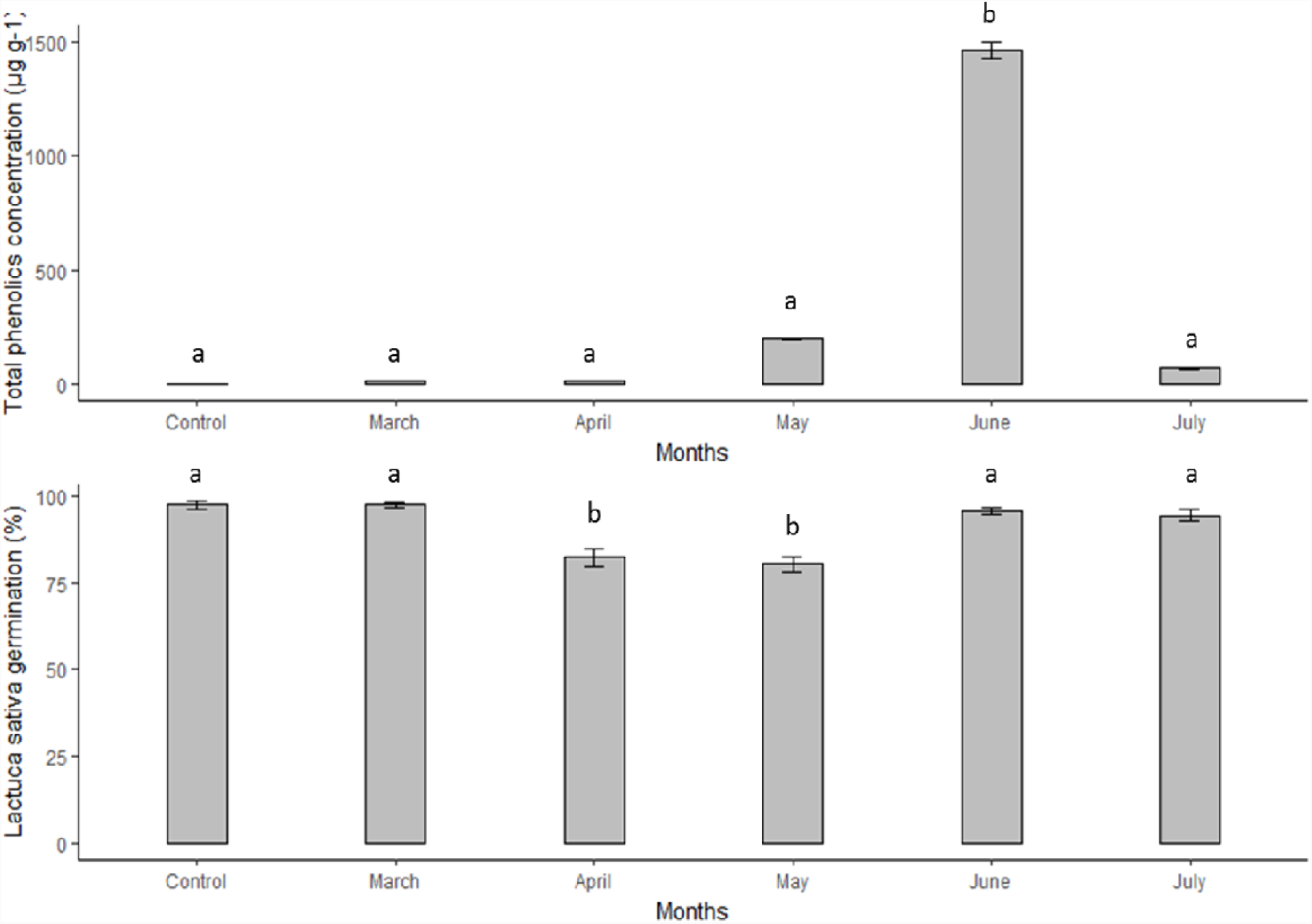
The monthly (March to July) total phenolic compound concentration (μg g^−1^) of the *R. ponticum* leaf leachate, and the mean germination percentage of the *L sativa* seedlings growing in the leachate. The error bars represent the standard error of the mean. Means with common letters did not differ significantly (P < 0.05) following one-way ANOVA and LSD analyses.

## DISCUSSION

Total phenolic compound concentration was significantly higher in June, contrary to our original hypothesis. Black et al. (2011) found seasonal variation in total phenolic compound concentration in the related species *R. tomentosum* to be closely correlated to day length in the Canadian arctic (Black et al. 2011). Whilst total phenolic compound concentration in our study coincided with the longest day length in June, the significant decrease in concentration from June to July suggests that day length was not solely responsible for the seasonal variation observed.

Other functions these compounds have within plants could also be responsible, including herbivore defence (Bennett and Wallsgrove 1994). Previous studies have shown young leaves and reproductive tissues have higher concentrations of secondary metabolites such as phenolics compared to older leaves, as they lack physical strength and are thus vulnerable to herbivory (Bennett and Wallsgrove 1994). This implicates that phenolic compound concentration should have been highest in March when the leaves were young, decreasing as the leaves gain physical strength as they age. Again however, our results contrast this, with total concentration was highest in June.

Despite total phenolics concentration being highest in June, *L sativ a* germination inhibition was highest during the spring months, consistent with our second hypothesis. Only quercetin was found to be significantly higher during one of these months however (May), a compound which was observed by de Martino et al. (2012) to inhibit the germination of garden cress. Other putative phytotoxins were also detected during May, including catechin, gallic acid and dihydromyricetin (Thorpe et al. 2009; Castro et al. 2010; Weidenhamer et al. 2013). However, these were present at higher concentrations in June (P < 0.05), and multiple regression analysis indicated that none were significantly correlated with suppressed germination.

These findings suggest that the observed germination suppression was not caused by an individual compound, rather it may be the result of complex interactions between compounds. The lack of a clear relationship between the seasonal variation in phenolic compound concentration and inhibition may also be influenced by the other roles phenolic compounds have within plants, as previously discussed.

The lack of understanding surrounding the roles of phytotoxic compounds in plant ecology is partly due to the complexity involved in designing an experiment which isolates the effect of the compounds from other environmental factors. Previous studies have used activated carbon which is reported to adsorb these compounds, however the addition of activated carbon may confound results by altering soil chemical properties (Weißhuhn and Prati 2009). Complexity is further increased by potential interactions with soil microbes, which are reported to mediate the effect of phytotoxic compounds, however this was not observed by Duke et al. (2009). We avoided using soil extracts in the current study, as the focus was purely on the synthesis of these compounds, in terms of both quantitative and qualitative seasonality, and the implications variations in these have on future studies in the field of chemical ecology. Whilst we acknowledge that soil interactions are not accounted for, we argue that the experimental design allowed the isolation of the effect from compounds released by other plants, in addition to seasonal variation in soil chemistry and environmental conditions.

We suggest that phytotoxic leaf leachate is just one of several interacting mechanisms (including shading, deposition of a deep litter layer and root exudation of phytotoxic compounds) that cumulatively contribute to the competitiveness of *R. ponticum.* The timing of inhibition appears to coincide with the germination and early growth period for many competing native seedlings.

The importance of phytotoxins in plant invasions is questionable, yet any such conclusions should consider seasonal variation, initial bioactivity and persistence in the soil. For example, the role of the phytotoxin catechin released from roots of the North American invasive *Centaurea stoebe* has been a topic of debate (Duke et al. 2009). Conflicting findings of studies on catechin in C. *stoebe* soils highlight the importance of considering temporal variation when investigating phytotoxcity mechanisms in invasive species, as the rate of production and release of individual phytotoxic compounds may be seasonally dynamic. Additionally, the concentration reaching the soil or leached away would depend on the intensity of recent precipitation. Compound persistence in the soil would also be dictated by microbial breakdown and adsorption processes in the soil (Blair et al. 2006; Weidenhamer et al. 2013). Finally, individual compounds may not act in isolation and likely interact synergistically even at low concentrations.

## ACKNOWLEDGEMENTS

GU and DG-J are grateful to both y Coleg Cymraeg Cenedlaethol and IBERS for supporting his Ph.D project stipend.

## AUTHORS’ CONTRIBUTIONS

GJ, MT, DGJ, JS and AW conceived the ideas and developed the experimental design; RO identified suitable sites for sampling; GJ and MT collected and analysed the data; GJ and DGJ led the writing of the manuscript. All authors contributed critically towards the manuscript drafts and gave final approval for publication.

## SUPPLEMENTARY DATA

**Table 1S:**
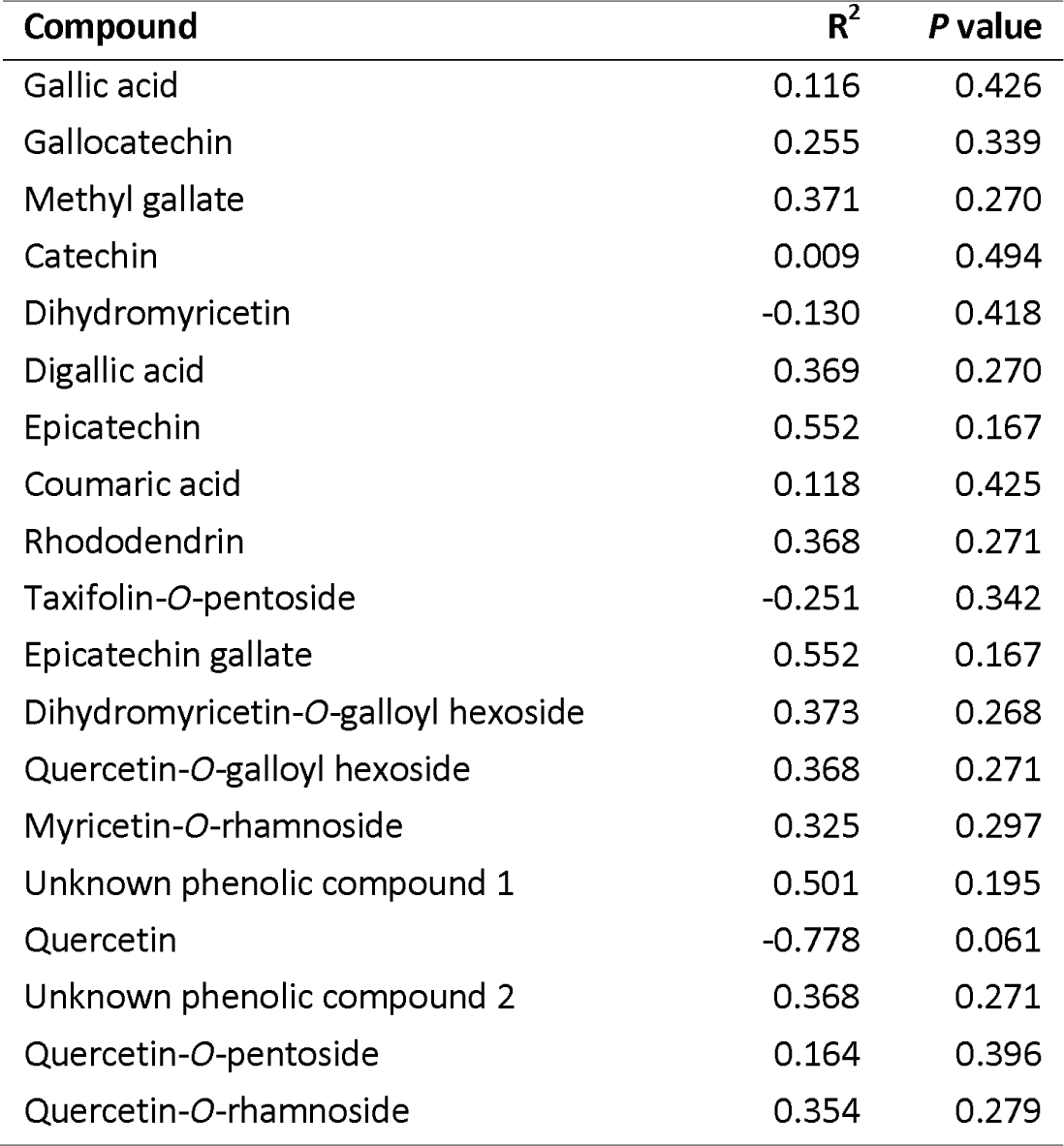
The R^2^ value and *P* value for each of the 19 compounds following a multiple regression analysis in the SPSS statistical software (significance level *P* < 0.05). Prior to analysis, the percentage germination data was arcsine transformed, whilst the concentration data for the compounds was log transformed.

**Table 2S:** Mean concentrations of the phenolic compounds detected in *R. ponticum* leaf leachate per tissue fresh weight (μg g^−1^) for each month. Total phenolics are also shown. Means with common letters were not significant different (P < 0.05) following One-way ANOVA; LSD Post Hoc Test.

| Compound and retention time (min) | March ($\mu\text{g g}^{-1}$) | April ($\mu\text{g g}^{-1}$) | May ($\mu\text{g g}^{-1}$) | June ($\mu\text{g g}^{-1}$) | July ($\mu\text{g g}^{-1}$) |
| --- | --- | --- | --- | --- | --- |
| Gallic acid (1.6) | $0.012 \pm 0.015$ a | $0.263 \pm 0.24$ a | $39.86 \pm 20.111$ a | $648.005 \pm 153.272$ b | $24.542 \pm 9.722$ a |
| Gallocatechin (3.4) | $0.01 \pm 0.01$ a | $0.087 \pm 0.087$ a | $0.316 \pm 0.205$ a | $5.106 \pm 5.097$ a | 0 |
| Methyl gallate (5.1) | $0.011 \pm 0.011$ a | 0 | 0 | $29.116 \pm 9.688$ b | 0 |
| Catechin (7.2) | $0.007 \pm 0.007$ a | 0 | $12.842 \pm 12.819$ a | $131.405 \pm 55.755$ b | 0 |
| Dihydromyricetin (9.3) | $0.001 \pm 0.001$ a | $0.029 \pm 0.019$ a | $14.979 \pm 9.386$ a | $37.922 \pm 3.507$ b | 0 |
| Digallic acid (11.6) | $0.007 \pm 0.007$ a | 0 | 0 | $10.905 \pm 5.071$ b | 0 |
| (Epi)catechin (12.7) | $0.062 \pm 0.062$ a | 0 | 0 | 0 | 0 |
| Coumaric acid (13.2) | 0 | $3.376 \pm 0.869$ a | $1.479 \pm 0.911$ a | $10.182 \pm 10.164$ a | $16.777 \pm 4.219$ a |
| Rhododendrin (14.9) | 0 | 0 | 0 | $46.936 \pm 18.49$ a | 0 |
| Taxifolin-*O*-pentoside (16.9) | $1.507 \pm 0.704$ a | $1.081 \pm 0.749$ a | $81.168 \pm 61.567$ a | $41.449 \pm 11.662$ a | $5.180 \pm 1.611$ a |
| (Epi)catechin gallate (17.6) | $7.510 \pm 7.497$ a | 0 | 0 | 0 | 0 |
| Dihydromyricetin-*O*-galloyl hexoside (18.3) | $0.009 \pm 0.009$ a | 0 | 0 | $1.958 \pm 1.955$ a | 0 |
| Quercetin-*O*-galloyl hexoside (22.9) | $0.001 \pm 0.001$ a | 0 | 0 | $8.125 \pm 8.11$ a | 0 |
| Myricetin-*O*-rhamnoside (23.6) | $0.067 \pm 0.061$ a | $0.198 \pm 0.197$ a | $2.684 \pm 1.581$ a | $211.43 \pm 70.467$ b | $5.501 \pm 3.223$ a |
| Unknown phenolic compound 1 (25.0) | 1.184 ± 0.277 a | 0.861 ± 0.531 a | 0.642 ± 0.403 a | 38.518 ± 12.9456 b | 2.612 ± 2.022 a |
| Quercetin (26.6) | 2.485 ± 1.232 a | 4.044 ± 0.901 a | 38.783 ± 19.01 b | 0 | 0 |
| Unknown phenolic compound 2 (27.6) | 0 | 0 | 0 | 3.144 ± 3.139 | 0 |
| Quercetin-*O*-pentoside (28.6) | 0.0.178 ± 0.162 a | 0.435 ± 0.162 a | 1.792 ± 0.434 a | 37.665 ± 6.768 b | 0 |
| Quercetin-*O*-rhamnoside (29.2) | 2.139 ± 0.523 a | 3.501 ± 0.331 a | 5.339 ± 1.855 a | 223.789 ± 80.68 b | 14.68 ± 7.618 a |
| **Total water-soluble phenolics** | 15.189 ± 0.395 a | 13.875 ± 0.293 a | 199.884 ± 4.623 a | 1466.241 ± 33.778 b | 69.292 ± 1.559 a |

